# From Proteins to Ligands: Decoding Deep Learning Methods for Binding Affinity Prediction

**DOI:** 10.1101/2023.08.01.551483

**Authors:** Rohan Gorantla, Ažbeta Kubincová, Andrea Y. Weiße, Antonia S. J. S. Mey

## Abstract

Accurate *in silico* prediction of protein-ligand binding affinity is important in the early stages of drug discovery. Deep learning-based methods exist but have yet to overtake more conventional methods such as giga-docking largely due to their lack of generalisability. To improve generalizability we need to understand what these models learn from input protein and ligand data. We systematically investigated a sequence-based deep learning framework to assess the impact of protein and ligand encodings on predicting binding affinities for commonly used kinase data sets. The role of proteins is studied using convolutional neural network-based encodings obtained from sequences and graph neural network-based encodings enriched with structural information from contact maps. Ligand-based encodings are generated from graph-neural networks. We test different ligand perturbations by randomizing node and edge properties. For proteins we make use of 3 different protein contact generation methods (AlphaFold2, Pconsc4, and ESM-1b) and compare these with a random control. Our investigation shows that protein encodings do not substantially impact the binding predictions, with no statistically significant difference in binding affinity for KIBA in the investigated metrics (concordance index, Pearson’s R Spearman’s Rank, and RMSE). Significant differences are seen for ligand encodings with random ligands and random ligand node properties, suggesting a much bigger reliance on ligand data for the learning tasks. Using different ways to combine protein and ligand encodings, did not show a significant change in performance.

**TOC Graphic:** 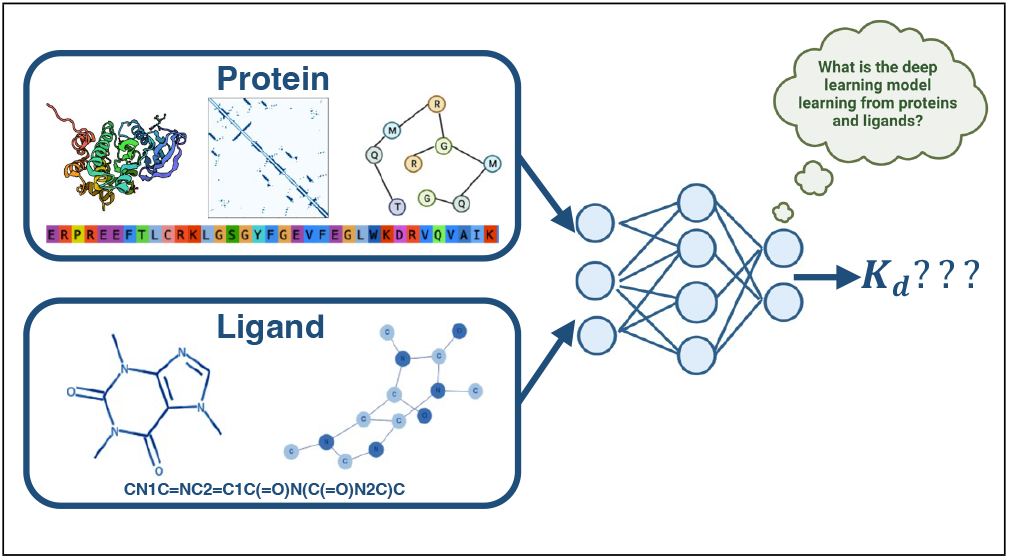

## Introduction

In computer-aided drug discovery, being able to predict the binding affinity (BA) between a protein and a potential drug candidate is critical to identify new small molecules from large libraries. Accurate experimental screening for good binders is not practical for rapidly testing millions of drug-like compounds against potential protein targets [1]. Over the last four decades, many different approaches to *in silico* predictions for binding affinities have been developed. This encompasses both structure-based and ligand-based approaches [2], however, each of them still has certain drawbacks when conducting a large-scale screening of compound libraries against a certain protein target. For example, docking [3, 4] methods can be used to screen large libraries, but often the desired accuracy for a BA is not achieved. On the other hand, alchemical free energy-based affinity prediction techniques [5–7] are more accurate, but computationally prohibitively costly for the discovery of hits in ultra-large libraries [8]. Both through the rapid development of new machine learning methods and better availability of binding affinity data, e.g. through PDBbind [9], KIBA [10], and Davis [11], many different efforts have been explored to generate ML-based methods for BA [12, 13].

In this paper, we will look at some of these machine learning (ML) models for binding affinity predictions more closely to gain insights on how components of these models contribute to the performance of the binding affinity prediction task. Depending on the type of input data used during training, these deep learning (DL) methods can be broadly categorized as sequence-or complex-based methods [2]. Complex-based methods [14–20], are trained on features from 3-dimensional (3D) protein-ligand complexes. These approaches typically use 3D convolutional neural networks (CNNs) or spatial graph neural networks (GNNs) to extract features from the 3D structure of protein-ligand complexes represented as 3D Cartesian grids, interaction fingerprints, and graphs. The extracted features of the protein-ligand complex are then used to predict the binding affinity.

Sequence-based approaches try to learn from Simplified Molecular Input Line Entry System (SMILES) strings and one-dimensional (1D) protein sequences. This can either be in the form of language models [21] or converting SMILES and protein sequences to graphs, leveraging 2D connectivity information from these graphs [22, 23]. The 1D and 2D-based DL models extract the features from the sequence and SMILES string and the feature vector formed by concatenating encoded protein and ligand features is used to get to the binding affinity prediction (Figure 1). Zhao et al. compiled a comprehensive overview of deep learning-based protein-ligand interaction prediction ML-based methods [13], which provides a useful starting point. We will take a closer look at some of the examples from this review, as our investigations focus on DL architectures from these examples.

**Figure 1:**
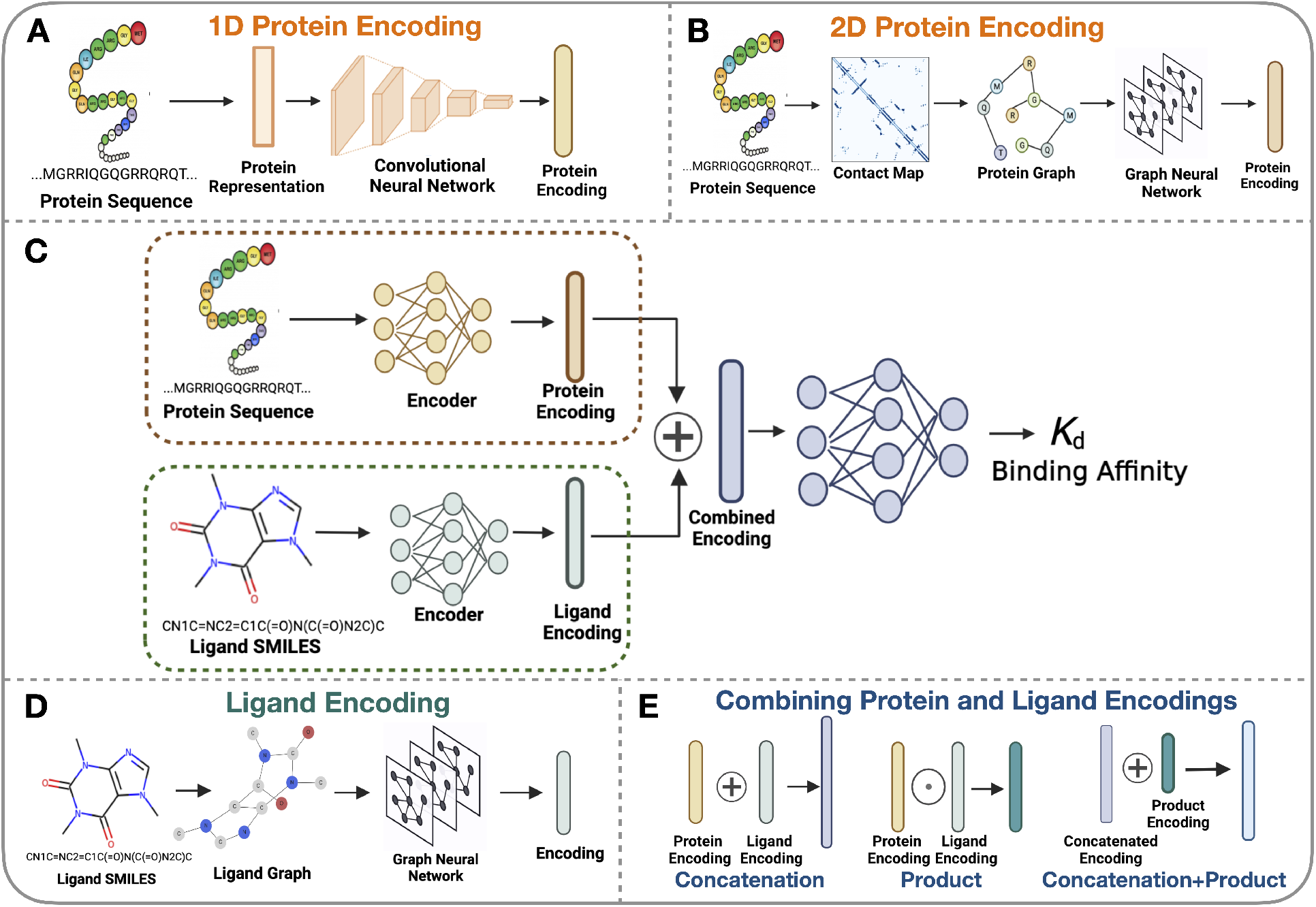
Systematic assessment of protein and ligand encodings on a deep learning framework for protein-ligand binding affinity predictions. **A:** 1D protein representation is obtained from the input sequence and then passed through a CNN module to obtain the protein-encoding. **B:** 2D protein encoding where an intermediary step of contact map prediction is required for the protein graph generation to obtain structural information from the sequences. The generated graphs are passed to graph neural networks to extract features and obtain the protein encodings. **C:** Overview structure of the DL framework used for this investigation. The DL framework processes the input sequence and SMILES data using 1D or 2D data structures to form their respective encodings. These encodings are combined and passed to a fully-connected neural network for binding affinity prediction. **D:** The input SMILES string is converted to a 2D graph and processed through the graph neural network to obtain the ligand encoding. **E:** Combination of protein and ligand encodings, namely concatenation, element-wise product, and concatenating the vectors from protein-ligand encoding concatenation and element-wise product.

Öztürk et al. [24] proposed DeepDTA, one of the earliest sequence-based methods using CNNs to extract 1D sequence information on the protein and ligand SMILES. WideDTA [25] extended DeepDTA by incorporating additional information sources, such as protein domains and motifs, and ligand maximum common substructure words. SMILES strings are a linearized representation of a ligand graph capturing structural, geometric, and topological properties.

Jiang et al. [23] introduced a more rational approach to utilize the information from the 2D contact map predicted by a supervised deep learning method, Pconsc4 [26], as the representation of the tertiary structure and have demonstrated improvements in binding affinity performance. These contact maps capture the details of residue-residue interactions and can be naturally modeled as graphs. All of these methods are trained and evaluated using publicly available kinase data sets [10, 11]. There are also other sequence-based DL methods [27–31] that have similar architectures to that of Jiang et al. [23].

More recently, Volkov et al. [32] highlighted challenges with current complex-based models, namely, that they do not necessarily learn the physics of protein-ligand binding. They found that explicit description of protein-ligand interactions from complexes provides no clear advantage compared to the corresponding interaction-agnostic models based solely on ligand or protein descriptors. Furthermore, Volkov et al. [32] discussed hidden protein and ligand biases in the PDBbind [9] data set for training complex-based models showing that these models have partly memorized the input data and did not learn the features that correspond to protein-ligand interactions.

In this paper, we open up the black box of typical CNN and G(C)NN architectures presented in the literature and investigate systematically how these model architectures learn from information presented to them through different encodings. We test ligand and protein encodings in 1D and 2D as summarized in Figure 1. For protein encodings, we look at 1D encodings obtained from sequences (Figure 1A) and 2D protein encodings obtained from contact maps (Figure 1B). For the 1D encodings, we compare the ESM1-b language model [33] to the performance of handcrafted KLIFS data using a one-hot encoding of the identified binding sites [34] on the downstream binding affinity prediction task. To test 2D encodings that rely on contact maps we use four different contact map prediction methods: protein sequence [33], homology information derived from multiple sequence alignment [26], and 3D structures [35] predicted through AlphaFold2. Lastly, we use a random contact map as a control. To study the impact of ligands on the DL framework, the input SMILES string is transformed into a graph structure and then processed using a GNN to obtain its encodings, as shown in Figure 1D. By looking at various perturbations of the ligand graphs, we can evaluate the effect on the downstream binding affinity prediction task. The last point of investigation is how the ligand and protein encodings are concatenated and how this may affect any binding affinity prediction Figure 1E. All experiments are carried out on the KIBA and Davis datasets as outlined in the methods section. Overall, similar conclusions to those of Volkov et al. were found that current architectures do not make much use of the protein data shown in these typical CNN and GCN architectures as presented in the results and discussion section.

## Methods

### Data sets

We used two kinase data sets, Davis [11] and Kinase inhibitor bioactivity (KIBA) [10], which are common benchmark data sets for the evaluation of how well deep learning (DL) models perform at binding affinity prediction tasks. Davis comprises of selectivity assays of 442 kinases and 68 inhibitors, with measurements for the inhibitor’s dissociation constants *K*_d_. Smaller *K*_d_ values mean higher affinity. These values were transformed into logarithmic space, consistent with prior studies [23, 24]. From Figure 2 B, it can be seen that there is a skew towards non-binders in this data set.

**Figure 2:**
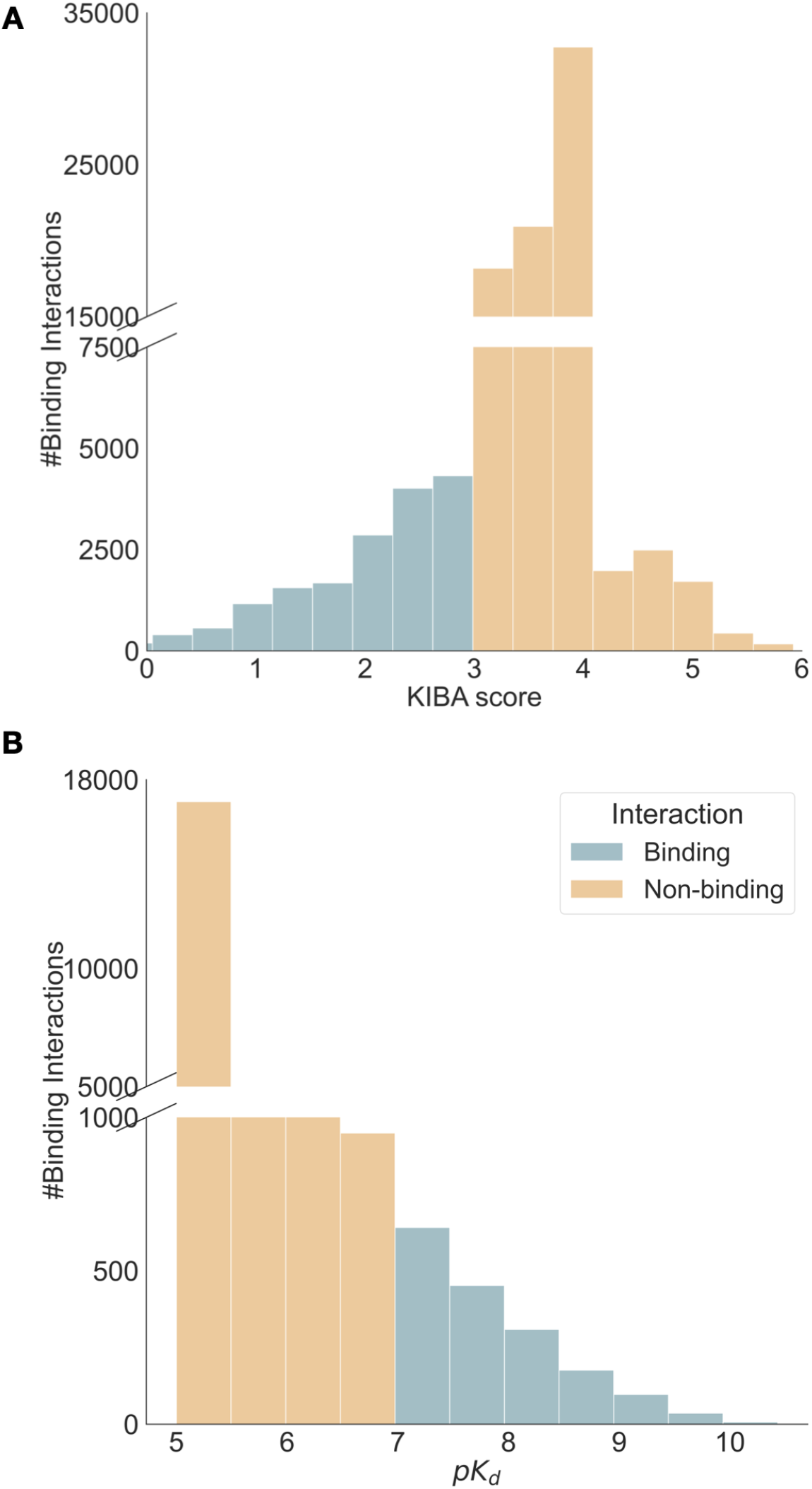
Summary statistics of KIBA[10] and Davis [11] data sets. KIBA has 188 proteins, 2111 ligands, and 95577 binding interactions, while Davis contains 333 proteins, 68 ligands, and 22644 binding interactions. **A:** Distribution of KIBA score across the entire data set. Lower KIBA score denotes higher binding affinity (*<*= 3) **B:** Distribution of *pK*_d_ scores across the Davis data set. Higher *pK*_d_ indicates a higher binding affinity with (*>* 7) usually seen as a binder.

The other data set, KIBA, amalgamates various sources of bioactivity data into a single KIBA score, optimizing consistency across different measures (*K*_i_*, K*_d_, and IC_50_), with lower scores implying a stronger binding affinity. The KIBA data set was originally comprised of 467 targets and 52,498 small molecules, however, He et al. [36] filtered it to contain only small molecules and targets with at least 10 observations yielding a total of 229 unique proteins and 2111 unique small molecules. This filtered data set was used to benchmark earlier BA prediction methods [22–24]. We filtered both the Davis and KIBA data sets further, to only include kinases with sequence lengths less than or equal to 1024 residues, as some of the protein-encoding techniques we used are limited to sequence sizes of up to 1024. Figure 2 summarizes the distribution and properties of both the Davis and KIBA data sets. Similarly to previous studies [22–24], we randomly divided each data set into six roughly equal parts, using 5*/*6 for training and validation, and the remaining data for testing.

### From features to encodings for ligands and proteins

To assess the performance of the BA learning task, we tested different encodings for ligands and proteins. We used graph-based approaches for ligands, and for proteins, we used 1D features and 2D graph-based encodings. In the following, we will outline these in detail.

#### Protein representations for generating 1D encodings

We tested two protein representations to study the effect of 1D encodings, Kinase–Ligand Interaction Fingerprints and Structures (KLIFS) and Evolutionary Scale Modeling (ESM-1b).

ESM-1b [33] is a protein language model based on a Transformer-34 [37] architecture trained on more than 220 million (unaligned) sequences from UniProt [38] through masked language modeling objective. During training, the transformer model is presented with protein sequences where a subset of their residues are masked, either by a random permutation to a different amino acid, by leaving them unmodified, or through a fraction of residues being masked. The objective of the model is to predict the values of the masked residues by considering the context of all unmasked residues in the input. We implemented the ESM-1b [33] model using the fair-esm Python package and the representations were obtained using esm1b t33 650M UR50S() model.

KLIFS provides information on how kinase inhibitors interact with their targets [34]. It provides a consistent alignment of 85 kinase ligand binding site residues that enables the identification of family-specific interaction features and the classification of ligands according to their binding modes. We leverage the 85 kinase ligand binding site residues for each kinase in the data sets using either Gene Name or UniprotID query. The 85 residues obtained for each kinase are then one-hot encoded; each residue is encoded to one of the twenty amino acids or a gap. The feature vector obtained from either KLIFS or the ESM-1b model is used as input to the convolutional neural network (CNN) module used for 1D encodings (see Figure 1A).

#### Contact maps for protein graph generation and 2D encodings

Protein’s 2D encodings are generated by means of a protein contact map. A protein contact map is a graph representation of a protein with an adjacency matrix containing information which amino acids in the protein chain are in contact or not [39, 40]. Protein graphs *G*_p_ = (**N**_p_, **M**_p_) are generated from the contact maps using input protein sequence with *L*_p_ residues. A pair of residues is said to be in contact or linked whenever the Euclidean distance (*d*_i,j_) between their C*_α_* atoms is less than or equal to a threshold *d*_c_. These connections can be determined from a 3D structure of a protein or predicted by means of other computational methods.

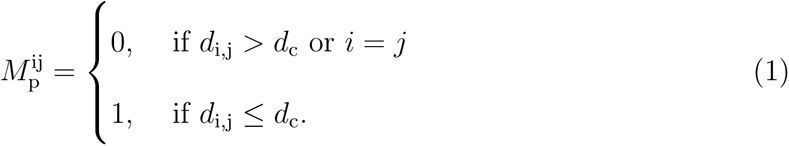

Equation 1, summarises the entries of the adjacency matrix **M**_p_ of the protein. In addition, each node in the adjacency matrix has certain node features which are represented by a matrix **N**_p_ *∈* ℝ*^L^*^p^*^×^*^54^. The 54-dimensional feature vector for each residue node is computed, for each of the *L*p amino acids with a summary of computed features presented in Table S1.

While the node features in the protein graph are kept constant, the contact map, i.e. the underlying adjacency matrix, is computed using four different ways to evaluate this. The first contact map prediction method used to obtain a protein graph is Pconsc4 [26]. It is a supervised DL method with a U-net architecture [41] trained on a curated data set with 2791 proteins from PDB and benchmarked on two data sets without homology to the training set [26]. It uses a 72-dimensional feature vector computed from multiple sequence alignment as input. The output of Pconsc4 is the probability of whether there is a contact between two pairs of amino acids, then a threshold of 0.5 is set to obtain the contact map, as proposed in the original paper [26]. The final contact map has a shape of (*L*_p_ *× L*_p_), where *L*_p_ is the number of nodes (residues or amino acids). This method was originally used by Jiang et al. [23] in their DL framework for binding affinity prediction.

The next method for obtaining a contact map is using data from an AlphaFold2 structural [35] model. AlphaFold2 predicts the 3D coordinates of all heavy atoms for a given protein using the amino acid sequence and aligned sequences of homologs as inputs. Here, we used the AlphaFold2 protein structure database [42] to get the 3D structures for each of the proteins used in the KIBA and Davis data sets. We downloaded the 3D structures in PDB format and used MDAnalysis version 2.0.0 [43] and NetworkX version 2.8.4 [44] to compute the contact map from the 3D structure. The pairwise C*_α_* distances were calculated for each given structure, and two residues were said to be in contact if their distance was less than 8 Å [39, 40].

We also used ESM-1b [33], as discussed earlier for extracting 1D protein encodings to obtain a contact map. The ESM-1b model predicts the contacts between residue pairs from the input protein sequence only. It learns the tertiary structure of a protein sequence in its attention maps during the unsupervised training on UniProt [38] data. The contact map predictions were made using esm1b t33 650M UR50S() model by calling model.predict contacts() method. At the time of our data collection for this study, ESM-1b was the most recent model available. However, it is worth mentioning that it has since been superseded by the release of the ESM-2 [45].

Randomly generated contact maps are used as a control method for studying the effect various contact map methods will have on protein graph encodings and, in turn, the binding affinity prediction. To generate random contact maps, we first generate a random protein sequence with randomly selected amino acid residues of the same length as the input protein sequence. The random sequence string is then used to get residue-residue contacts using the ESM-1b [33] model in a similar way as described above. Using the contact information from each of discussed methods, we then compute the adjacency matrix **M**_p_ to build the protein graph.

#### Ligand encodings and their perturbations

The ligands are represented as graphs derived from a linearised version of their chemical structure represented as SMILES strings. Ligand encodings are then obtained from these graph representations. Ligand graphs *G*_l_ = (**N**_l_, **M**_l_) are generated from the input SMILES string with *L*_l_ atoms, where **M**_l_ *∈* ℝ*^L^*^l^*^×L^*^l^ is an adjacency matrix with information about the chemical bonds present between any given pair of atoms. Self-loops are added to the graph construction, i.e., the diagonal of the adjacency matrix is set to one to improve the feature performance of the molecule. Each ligand node in the graph is denoted by a 78-dimensional feature vector similar to Jiang et al. [23] capturing the one-hot encoding of the atom type, degree of the atom, total number of hydrogens bound to the atom, number of implicit hydrogens bound to the atom, and whether the atom is aromatic or not. We processed the SMILES string with RDKit version 2020.09.5 [46] library using Chem.MolFromSmiles() to get the atom and bond details for building the ligand graph.

To look at the effects of the ligand encodings on the downstream task we designed three different ways to perturb the ligand graphs. The first randomization technique, we call *Point randomization*, generates a new SMILES string with minor changes to the original one, thus altering the ligand graph slightly. In this process, we enumerate the presence of certain atoms (such as Cl, F, Br, and (***−***O)) within the string, and selectively modify up to four atoms. This can involve substituting one halogen atom with another or removing a (***−***O) atom. In cases where none of the enumerated atoms exist, a Cl atom is appended at the beginning of the SMILES string. More details about the point randomization algorithm are provided in the SI. This variation helps us ascertain if small changes in ligand structure can influence binding affinity prediction and identify if the model accurately captures these structural alterations.

The second technique, *Node feature randomization*, assesses the impact of node features in the model’s predictions. We randomly permute the node feature values across the graph, thus disrupting the nodes’ identities while preserving the graph’s structure. The degree of performance change following this randomization will indicate the model’s dependency on node features versus the graph structure for its predictions. The third method, *Random sampling*, represents an extreme level of randomization, where the original ligand graph is substituted with a randomly selected ligand graph from the same data set. This approach enables us to evaluate whether our DL model relies on ligand features for binding affinity prediction. By training the model with a randomly selected ligand, we can ensure that we are not generating chemically implausible ligands.

### Deep learning architecture

To combine information from our ligand encodings and protein encodings in a deep learning architecture, we borrow ideas from the architecture proposed by Jiang et al. [23]. We use a module for protein encodings and one for ligand encodings. For the 1D protein encodings, we use a three-layer CNN model (Figure S1). For the 2D graph approaches used for both ligand and protein a GNN model with three graph convolutional network (GCN) layers similar to Jiang et al. [23] is used (Figure S2). The GCN model learns the representation for a given input graph *G* = (**N**, **M**), where **N** *∈* ℝ^v^*^×^***^q^** is the matrix containing v nodes and each node is represented by a **q** dimensional feature vector. **M** *∈* ℝ^v^*^×^*^v^ is the adjacency matrix that provides the structural information of the graph. The features are extracted from the graph via GCN layers, where each layer will perform a convolution operation by following the propagation rule [47] defined below

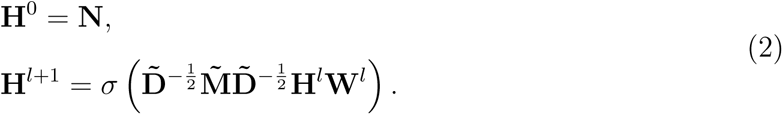

Here, **H***^l^* and **W***^l^* denote the *l*^th^ GCN layer outputs and its corresponding learnable parameters respectively. The adjacency matrix, 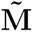 with self-loops in each node, i.e., 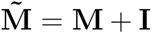, where **I** is the identity matrix and 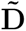 is the diagonal node degree matrix calculated from 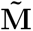, 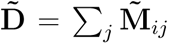 The design of the 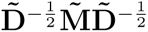 term is intended to add a self-connection to each node and keep the scale of the feature vectors. *σ* (.) represents a non-linear activation function, Rectified Linear Unit (ReLU).

### Experimental setup for model training and analysis

The detailed DL architecture is outlined in the SI. Figure S1 summarises the CNN architecture used for the 1D protein and ligand encodings with the 1D protein encoding making use of the highlighted CNN module. Figure S2 contains a summary of the architecture used for the GCN of the 2D protein-ligand encodings. These protein graphs *G*_p_ and ligand graph *G*_l_ derived encoding is obtained using the GCN module. Both the CNN and GCN modules are implemented with PyTorch and PyTorch geometric. We use the Mean Squared Error (MSE) loss function to train the DL model. Experiments testing combinations of ligand and protein encodings in 1D and 2D are summarized in Table S2. Each experiment trains the DL model for 2000 epochs with batch size 128 and learning rate *β* = 0.001 using the Adam optimizer, saving the top-performing model from the validation set. To ensure the robustness of our experiments, we randomly selected three deep learning models trained on three different folds from the training split. These models were then used for bootstrap resampling on a randomly selected sample size of 1500 data points from the test set. The mean predictions of the trained models for each bootstrap iteration are used to compute the evaluation metrics and their associated errors. We use Concordance Index (CI), Root Mean Squared Error (RMSE), Pearson correlation, and Spearman rank correlation to assess the model’s performance. All code and models are accessible at https://github.com/meyresearch/DL_protein_ligand_affinity.

## Results and discussion

### Different protein contact map prediction methods provide different protein graphs for protein encodings

We computed protein contact maps (PCM) for the KIBA and Davis data sets, as outlined in the methods section using structural data from AlphaFold2, the sequence-based method ESM-1b, and the homology modeling tool Pconsc4. To obtain a baseline idea of how well these methods correlate to experimentally determined structure-derived PCMs, we manually curated 50 protein structures with structural data available in the RCSB protein data bank (PDB) spanning across the kinase data sets KIBA and Davis. These PDB structures were identified from the PDB using either Uniprot ID or Gene ID. Contact maps from PDB data were computed in the same way as AlphaFold2 PCMs, but used as a reference. To evaluate the performance of contact map prediction methods, we used the F1 score, Matthews’ correlation coefficient (MCC) [48], and precision metrics. MCC is a balanced metric that considers the distribution of true positives, false positives, true negatives, and false negatives in a binary classification problem, making it a suitable metric for evaluating models in cases of class imbalance; we provide more details about the metric in the SI. Figure 3A shows an example of a PCM obtained from PTK-6 with Uniprot ID: Q13882 and PDB ID 5D7V with 8 Å threshold and highlight the true contacts (turquoise squares), falsely predicted contacts (pink circles) and lost contacts (orange crosses), that is, those that were present in the 3D X-ray structure contact map but not present in the predicted one. For more examples, see the SI.

**Figure 3:**
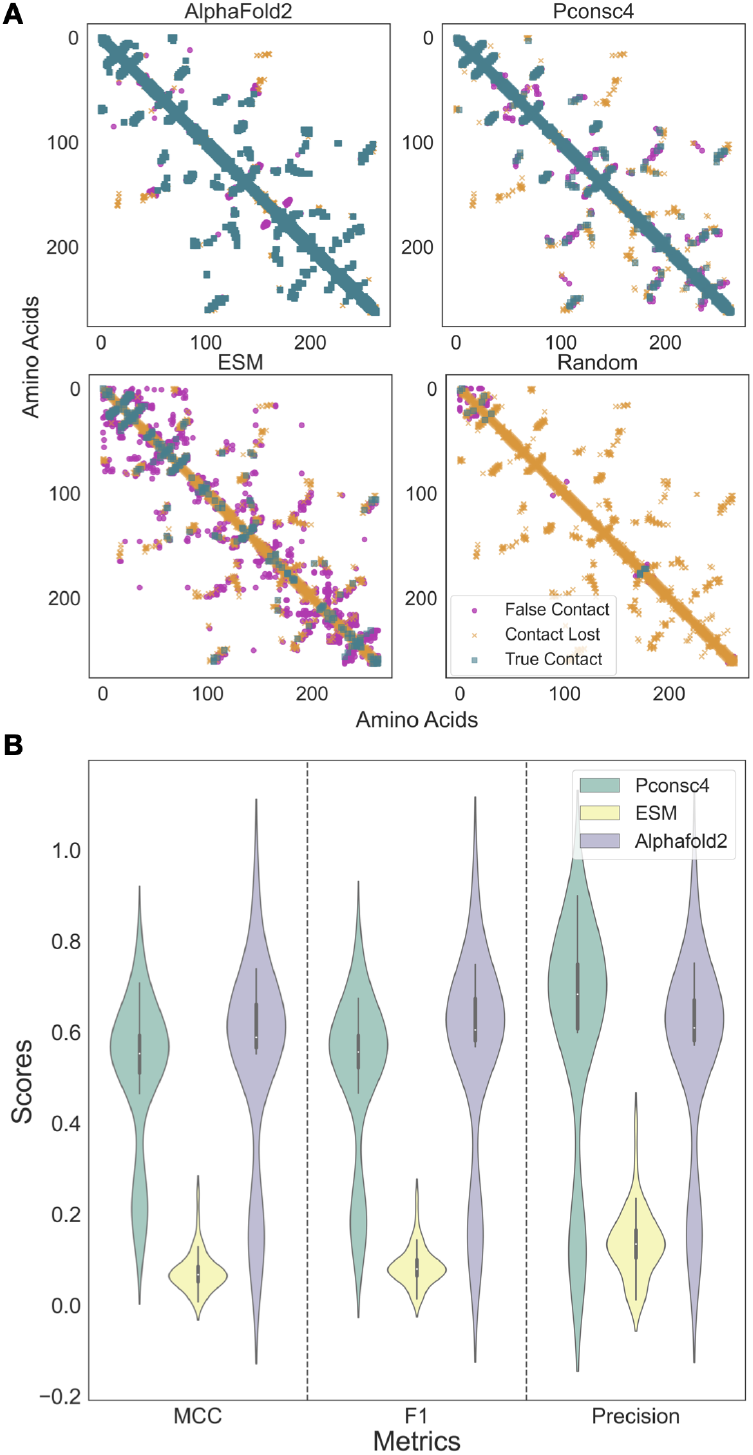
Contact maps obtained from each contact map prediction algorithm (ESM-1b, AlphaFold2, and Pconsc4) are different and provide significantly different protein graphs to the protein encodings. **A:** Contact map analysis for PTK-6 using PDB ID 5D7V as a reference, top left AlphFold2, top right Pconsc4, bottom left ESM-1b and bottom right a random contact map. True contacts are displayed in turquoise squares, lost contacts are shown in orange crosses, and falsely predicted contacts in pink circles. **B:** Assessing contact map methods on a curated data set of KIBA and Davis protein structures, show that the AlphaFold2 contact maps perform better on MCC, and F1 score metrics, while the Pconsc4 contact map prediction method has higher mean precision. ESM-1b contact predictions are the least reliable.

Figure 3B shows violin plots that compare PCM generation methods and their performance according to MCC, F1, and Precision. The contact map predictions from AlphaFold2 structures had the highest mean MCC and standard deviation of 0.54 *±* 0.21 and F1-score of 0.55 *±* 0.22, while the ESM-1b method had the lowest average MCC (0.07 *±* 0.04) and F1-score (0.09 *±* 0.04). The contact map prediction results of Pconsc4 (MCC-0.51 *±* 0.16, F1-score-0.51 *±* 0.17) are comparable to that of the AlphaFold2; however, the mean precision of Pconsc4 (0.59 *±* 0.25) method is slightly better than AlphaFold2 (0.55 *±* 0.22) contact maps on the curated PDB data set. Using the Wilcoxon signed-rank test, we evaluated the performance of the contact map prediction methods on the 50 experimentally determined structures. In the Wilcoxon signed-rank test, the null hypothesis is that there is no significant difference between the performance of the two models. Generally, a p-value less than or equal to the significance level of 0.01 is considered statistically significant, leading to the rejection of the null hypothesis. On comparing the method predictions on MCC, F1-score and precision for each pair of methods we observed a small p-value (*p <* 0.01) indicates strong evidence against the null hypothesis. Thus, there is a significant difference between each pair of the contact map prediction methods used in this study, and the contact map predictions obtained from each method are not the same. ESM-1b appears to be the least reliable and accurate method for predicting contact maps, whereas AlphaFold2 and Pconsc4 exhibit almost biomodal distributions for MCC and F1-score. To understand the reliability of protein structures used to obtain AlphaFold2 contact maps, we computed the average confidence score of AlphaFold2 structures per residue in the Davis and KIBA data sets. The average confidence score per residue for KIBA was 78.2 *±* 21.52, while for Davis, the score was slightly higher at 88.82 *±* 22.35. Removing low confidence score (*<* 70) AlphaFold2 structures from the set of 50 hand-curated X-ray structures does not remove the bimodal distribution in MCC, F1, and Precision for AlphaFold2 and Pconsc4 and other underlying factors beyond the scope of this paper give rise to this.

### Protein encodings based on significantly different protein graphs do not have much effect on binding affinity prediction

With four different ways to generate protein graphs established, we want to assess if the protein graph structure has any impact on the downstream binding affinity prediction task when we use the PCM in our protein encoding. We will refer to these as 2D encodings as we are generating graphs with nodes and edges and interpreting them as 2D structures.

Keeping the deep-learning framework fixed, as described in the methods and shown in Figure 1C, we only test the four different contact map generation methods. The ligand encodings are untouched and based on the DL-framework from [23]. The downstream task of estimating binding affinities is evaluated on both the KIBA and Davis data set.

Figure 4 A (KIBA) and B (Davis) summarise the findings of changing the PCM generation methods. From Figure 4 A and 4B, we can observe that there is not much change in the performance of the DL model with different protein encodings across all four evaluation metrics, i.e., CI, Pearson correlation coefficient, RMSE, and Spearman rank correlation on the test set. The experiments on the Davis data set in Figure 3D show that the random encoding (CI: 0.86 *±* 0.01, Pearson: 0.79 *±* 0.02, RMSE: 0.51 *±* 0.02) has slightly lower performance than Pconsc4 (CI: 0.89 *±* 0.01, Pearson: 0.82 *±* 0.01, RMSE: 0.48 *±* 0.02), ESM-1b (CI: 0.89 *±* 0.01, Pearson: 0.82 *±* 0.01, RMSE: 0.47 *±* 0.02), and AlphaFold2 (CI: 0.88 *±* 0.01, Pearson: 0.82 *±* 0.02, RMSE: 0.49 *±* 0.02), while there is no change among the rest of the methods. On the KIBA data set, ESM-1b has the lowest Root Mean Squared Error (RMSE) on the test set, with 0.468 *±* 0.02, followed by Pconsc4 with 0.475 *±* 0.02, and Random and AlphaFold2 with 0.480 *±* 0.03. Pearson’s correlation between experimental and predicted binding affinity score for ESM-1b and Pconsc4 was 0.82 *±* 0.02, while Random and AlphaFold2 had 0.81 *±* 0.02. From both Davis and KIBA data sets, we can observe that the Random encoding performed comparatively worse than the rest.

**Figure 4:**
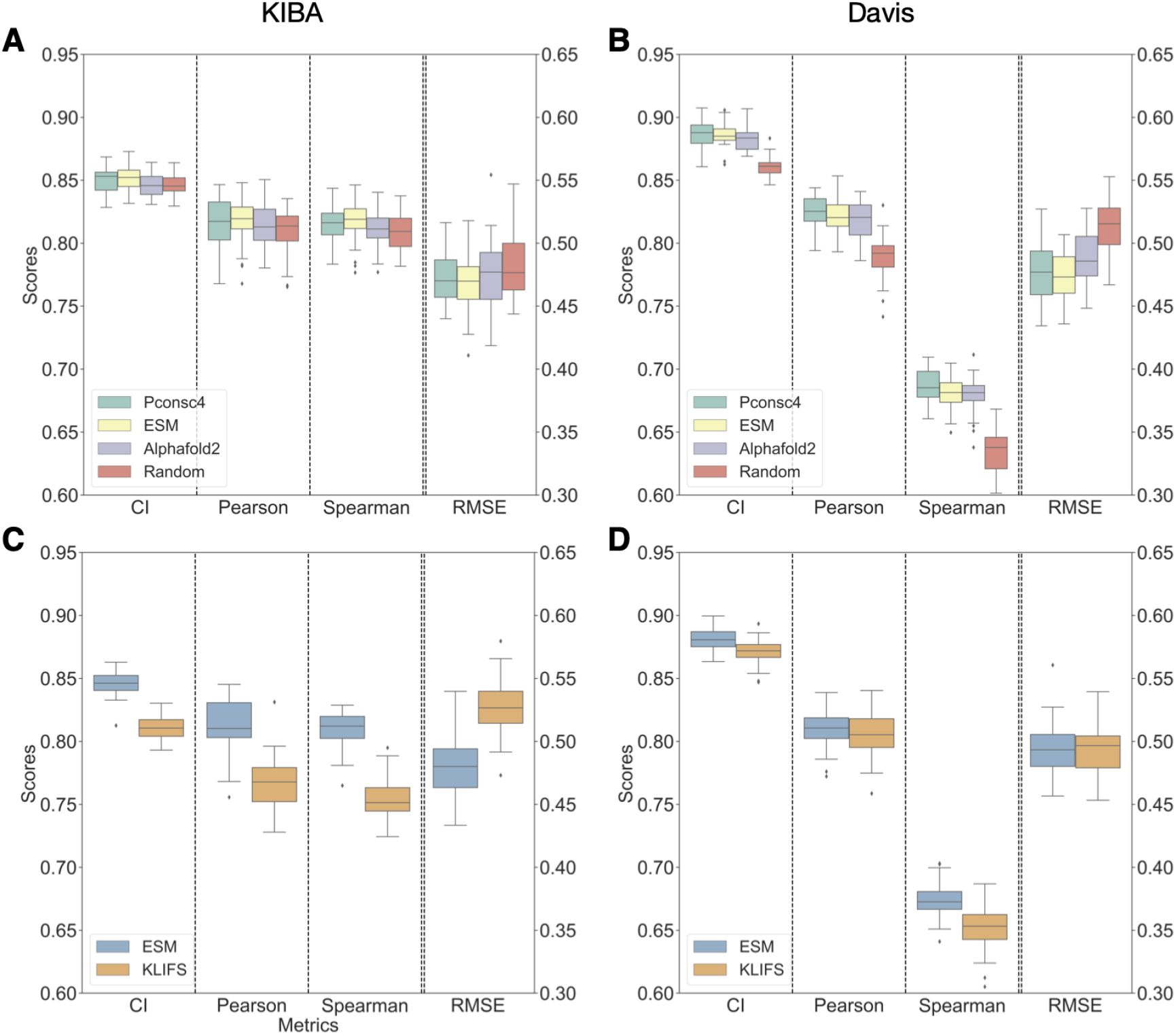
Protein encodings (2D) with structural information from contact maps do not have much effect on binding affinity prediction and 1D encodings from protein language models (PLM) perform similarly to contact maps enabled encodings. **A, B:** Boxplots for different performance measures (CI, Pearson correlation, Spearman Rank, and root mean square error) of binding affinity predictions for the KIBA data set (A) and Davis data set (B) for four different protein contact map methods. This shows that the structural information from protein contact maps encoded into a graph is not making any significant contribution to DL model performance. **C, D:** Boxplots for two different 1D encoding methods and their performance metrics (CI, Pearson correlation, Spearman Rank, and root mean square error), the PLM encodings of the ESM-1b model perform better than one-hot encodings from KLIFS handcrafted sequences on both the data sets. Overall, the performance of 1D encodings is comparable to the encodings that include information from protein graphs.

Using the Wilcoxon signed-rank test, we evaluated the performance of the trained DL models on the bootstrapped test set. The Wilcoxon signed-rank test evaluates whether there’s a meaningful difference between two models’ performances and a p-value of 0.01 or less generally suggests this difference is statistically significant, refuting the original assumption of no difference. The KIBA data set shows no significant difference (*p >* 0.01) in the performance of AlphaFold2, ESM-1b, Pconsc4, and Random models in terms of Pearson, Spearman, and RMSE metrics. The overall performance of all metrics is better on the Davis data set, with no significant difference (*p >* 0.01) between AlphaFold2 and Pconsc4. However, ESM-1b has a significantly better performance on all metrics (*p <* 0.01) on the Davis data set. We also observe a significant drop in performance with random encodings on the Davis data set. This performance drop is higher than for KIBA as both these data sets have different proportions of proteins and ligands (Davis-334 kinases and 68 ligands, KIBA-188 kinases and 2111 ligands).

Next, we looked at the overall correlation of either KIBA score or *pK_D_* predictions for each molecule in the test set. We arbitrarily picked Pconsc4 as a reference to compare the predictions with various encoding methods (Figure S5). Each pair of methods exhibits a strong correlation in their binding affinity predictions. All the pairs compared had a coefficient of determination, *R*^2^, that was close to 1, indicating high comparability in their predictions. Figure S6 shows the correlation between experimental and predicted binding affinities along with the kernel density estimate (KDE) of the prediction distributions for all four 2D encoding methods. For the KIBA data set, the *R*^2^ for all the methods is close to 0.71 *±* 0.01, while for Davis the *R*^2^ for ESM-1b, Pconsc4, and AlphaFold2 is 0.72 *±* 0.01 and the *R*^2^ for random is 0.69 *±* 0.02. We compared the prediction distributions of each pair of methods for both KIBA and Davis using the Jensen-Shannon (JS) divergence. JS divergence is a symmetric measure that quantifies the similarity between two probability distributions. It is a smoothed version of the Kullback-Leibler (KL) divergence and is bounded between 0 and 1, with 0 indicating identical distributions and 1 indicating completely different distributions. The mean JS divergence for all pairs of methods on KIBA is 0.0008, and for Davis, it is 0.0031. This shows that the prediction distributions from each method are similar to one another, and there is not much change in model predictions with encoding generated from significantly different contact maps. Overall, these results suggest that there is no significant change across AlphaFold2, Pconsc4, and ESM-1b on the data sets, while the DL model trained with random contact maps showed a slight drop in performance on the Davis data set.

### Encodings from protein language models outperform handcrafted encodings in predicting binding affinity and perform similarly to 2D protein encodings

Given that different contact maps, including random maps show little impact on the accuracy of the binding affinity prediction. Next, we use 1D encoding methods to investigate the importance of structural details in the DL model’s prediction capabilities. To this end, we use 1D encodings obtained from ESM-1b, and handcrafted sequences that identify binding regions explicitly as contained in KLIFS sequences [34]. If we show the model only part of the sequence, but that part is known to be part of the binding site of the protein, will this improve learning? The results that address these questions are presented in detail in Figure 4 C and D. Figure S7 shows a Euclidean distance heatmap of the ESM-1b embeddings capturing variance among the proteins in both KIBA and Davis data sets.

Figure 4C and 4D show that the ESM-1b encodings-based model performs better than the one-hot encoding using KLIFS sequences for both KIBA and Davis data sets. For KIBA, the ESM-1b-based encoding (CI: 0.84 *±* 0.01, Pearson: 0.81 *±* 0.01, RMSE: 0.48 *±* 0.02) is performing better than KLIFS-based one-hot encodings (CI: 0.81*±*0.01, Pearson: 0.77*±*0.01, RMSE: 0.51 *±* 0.02). The Wilcoxon signed-rank test on both these encodings on the KIBA data set shows that the change in performance is significant (*p <* 0.01) across all the metrics. From Figure 4D, the performance of both ESM-1b (CI: 0.88 *±* 0.01, Pearson: 0.81 *±* 0.01, RMSE: 0.49 *±* 0.02) and KLIFS (CI: 0.87 *±* 0.01, Pearson: 0.81 *±* 0.01, RMSE: 0.49 *±* 0.02) the Davis data set is comparable. The Wilcoxon test shows that the change in performance is significant on CI and Spearman metrics with *p <* 0.01, while on Pearson and RMSE, the change is not significant (*p >* 0.01). We can see that both the rank correlation metrics CI and Spearman have seen a significant performance change between PLM encodings and manually curated sequence-based encodings. Further, from our 1D encoding experiments, we can see that 1D ESM-1b encodings perform similarly to the 2D encodings on both KIBA and Davis data sets (Figure 4A and 4B) with no significant change *p >* 0.01 in performance with 2D ESM-1b and Pconsc4 encodings. This shows that adding structural information in the form of a protein graph based on a contact map did not improve the overall performance of the DL model significantly.

### The deep learning model relies on good ligand encodings for learning binding affinities

Now we make changes to the ligand encodings as laid out in the methods section to systematically assess how ligand encodings contribute to the overall learning task. From 5A and 5B we can see that the model’s performance on the test set that the ligand encodings greatly impact the binding predictions on both data sets. On both KIBA and Davis data sets, the point randomized encoding had the lowest drop in performance as compared to random node and random sampling perturbation methods. For point randomization methods, there is less than 1% drop across all metrics on KIBA, whereas on Davis there is 3.65% *±* 1% on CI and 8.11% *±* 2% on Pearson metrics. However, the Wilcoxon test for point randomization perturbation as compared to the original ligand encoding has *p <* 0.01 on both data sets, denoting the change to be significant. In randomizing node feature perturbation, there is a drastic drop in performance on both data sets. The performance on KIBA dropped by 17.18% *±* 2% on CI, 45.82% *±* 4% on Pearson, and the RMSE increased by 86.03% *±* 0.6%. For Davis, the changes are even more drastic with 34.78% *±* 1% on CI, 73.09% *±* 3% on Pearson, and the RMSE increased by 90.07% *±* 4%. The performance of random sampling perturbation has a similar stark effect as node feature randomization for both KIBA (CI: 27.15% *±* 1%, Pearson: 82% *±* 2%) and Davis (CI: 37.96% *±* 1%, Pearson: 80.81% *±* 3%). Both random node and random sample perturbations have a significant change (*p <* 0.01) in binding affinity performance as compared to the original ligand encoding. We can observe that the performance for Davis dropped more than for KIBA; this could be due to the difference in data set distributions, as the number of ligands in Davis (68) is much smaller than for KIBA (2111).

In the SI we also highlight what effect the changes on ligand encodings have in terms of actually predicting binding affinities with respect to experimentally observed values. The findings are summarized in Figure S8. Original ligand encodings obtained *R*^2^ on the test set for KIBA 0.71 *±* 0.01 and 0.72 *±* 0.01 for Davis. Point randomizations have a more distinct effect for Davis with a drop of *R*^2^ = 0.64 *±* 0.01, compared to KIBA *R*^2^ = 0.70 *±* 0.01. One possible explanation for this is the smaller ligand data set size for Davis, and another is the KIBA score choice itself, which will be discussed in more detail below. For the random node encodings, *R*^2^ = 0.14 *±* 0.00 for KIBA, *R*^2^ = 0.01 *±* 0.03, for Davis, and similarly for the randomly sampled ligand encodings, *R*^2^ = 0.01 *±* 0.046 for KIBA and *R*^2^ = 0.02 *±* 0.13 for Davis. Introducing the randomizations in the ligand encodings the deep learning model can no longer perform the learning task and the resulting model is unusable as a potential model for binding affinity predictions for kinases. An obvious conclusion is that the presented architecture predominantly learns from ligand encodings and protein features play hardly any role.

### Combining protein and ligand encodings in different ways has no significant effect on the model’s predictability

Lastly, we look at how ligand and protein encodings can be combined in the DL framework. Jiang et al. [23] used concatenation operations to combine protein and ligand encodings with the combined vector being passed along the fully connected layers to predict binding affinity [22–24]. Other combination methods are possible: the element-wise product of protein and ligand encodings and the concatenated vector obtained by combining both element-wise product and concatenation operations. Concatenation allows the DL model to learn complex interactions between the protein and ligand features; here, the model will have access to all the features of the protein and ligand. On the other hand, the element-wise product emphasizes the DL model to learn features important for both protein and ligand. Here, we provide the model with a feature space that is expected to have the most informative aspects of the protein and ligand encodings. When the protein and ligand encoding is concatenated with the product encoding, the model will have access to a feature space that is larger and richer than that in either approach alone.

From Figure 6A (KIBA) and 6B (Davis), we can see no significant improvement in the performance of the DL model on the binding affinity prediction task by both element-wise product of protein and ligand encodings and the concatenated vector obtained by combining both element-wise product and concatenation operations. On the Davis data set, there is a slight drop in performance with the element-wise product (CI: 0.88 *±* 0.01, Pearson: 0.81 *±*, Spearman: 0.67*±*0.01) as compared to both the concatenation (CI: 0.89*±*0.01, Pearson: 0.82 *±* 0.01, Spearman: 0.69 *±* 0.01) and fusion encoding of concatenation and element-wise product (CI: 0.89 *±* 0.01, Pearson: 0.82 *±* 0.01, Spearman: 0.69 *±* 0.01). From the Wilcoxon test, the drop by incorporating the element-wise product in the place of concatenation is significant (*p <* 0.01). In contrast, the improvement with fusion concatenation and the element-wise product is not statistically significant (*p >* 0.01). From experiments on the KIBA data set, the element-wise product (CI: 0.85 *±* 0.01, Pearson: 0.81 *±* 0.02) encoding performed almost the same as the concatenation (CI: 0.85 *±* 0.01, Pearson: 0.81 *±* 0.02) and fusion encoding (CI: 0.85 *±* 0.01, Pearson: 0.82 *±* 0.02). The performance for the KIBA data set for both methods shows no statistically significant change (*p >* 0.01). The DL model is not learning anything new from the element-wise product and the fusion of concatenation and product feature spaces.

**Figure 5:**
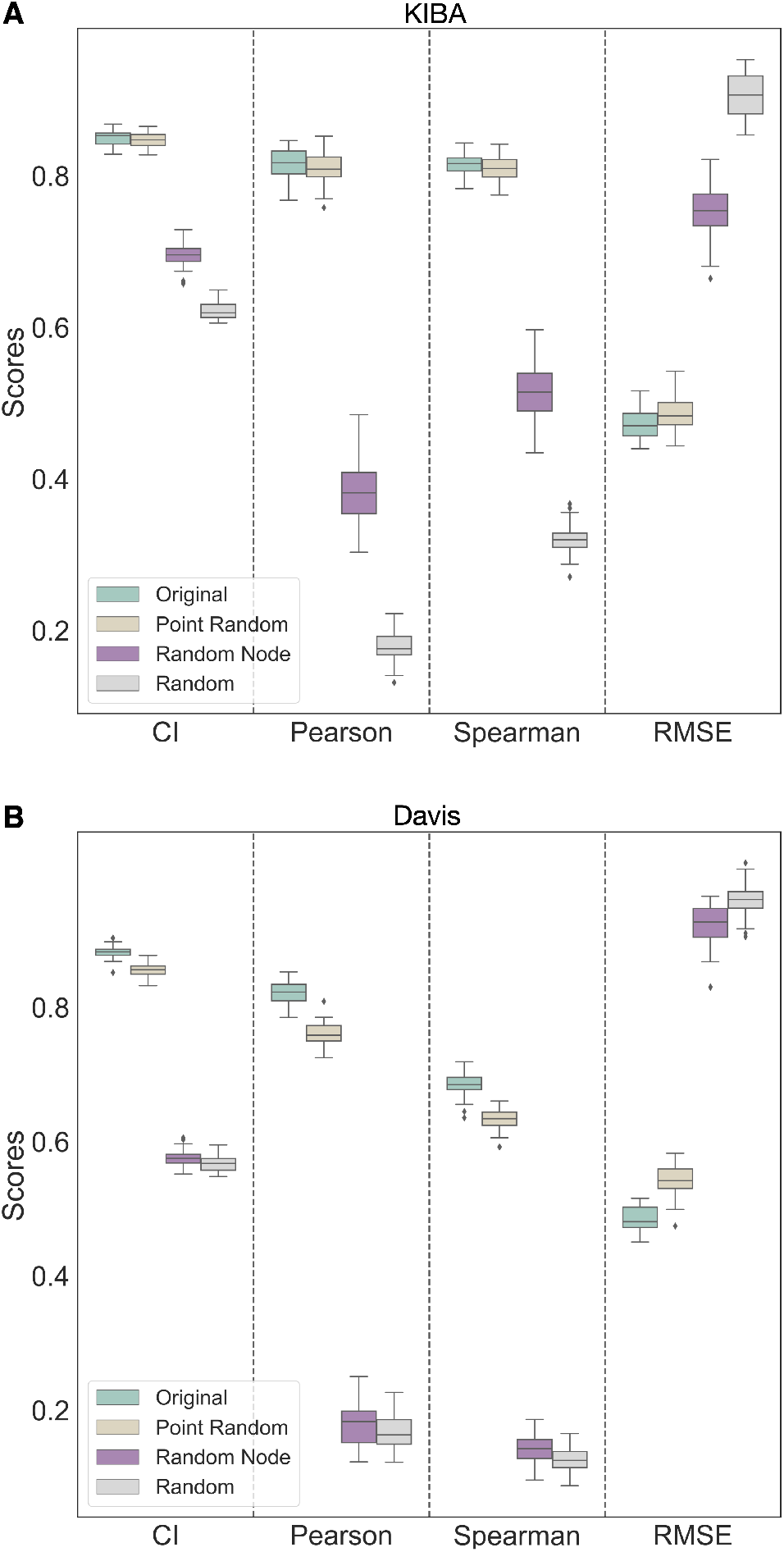
Changes to ligand encodings show a significant change in binding affinity performance. Comparative analysis of the DL model performance in binding prediction testing four different ligand encodings shows that the DL model relies on ligand encodings for both KIBA (**A**) and Davis (**B**) data sets. The DL model with randomly sampled and randomized node feature encodings are not learning to estimate BA during training, demonstrating the model’s reliance on ligand information.

**Figure 6:**
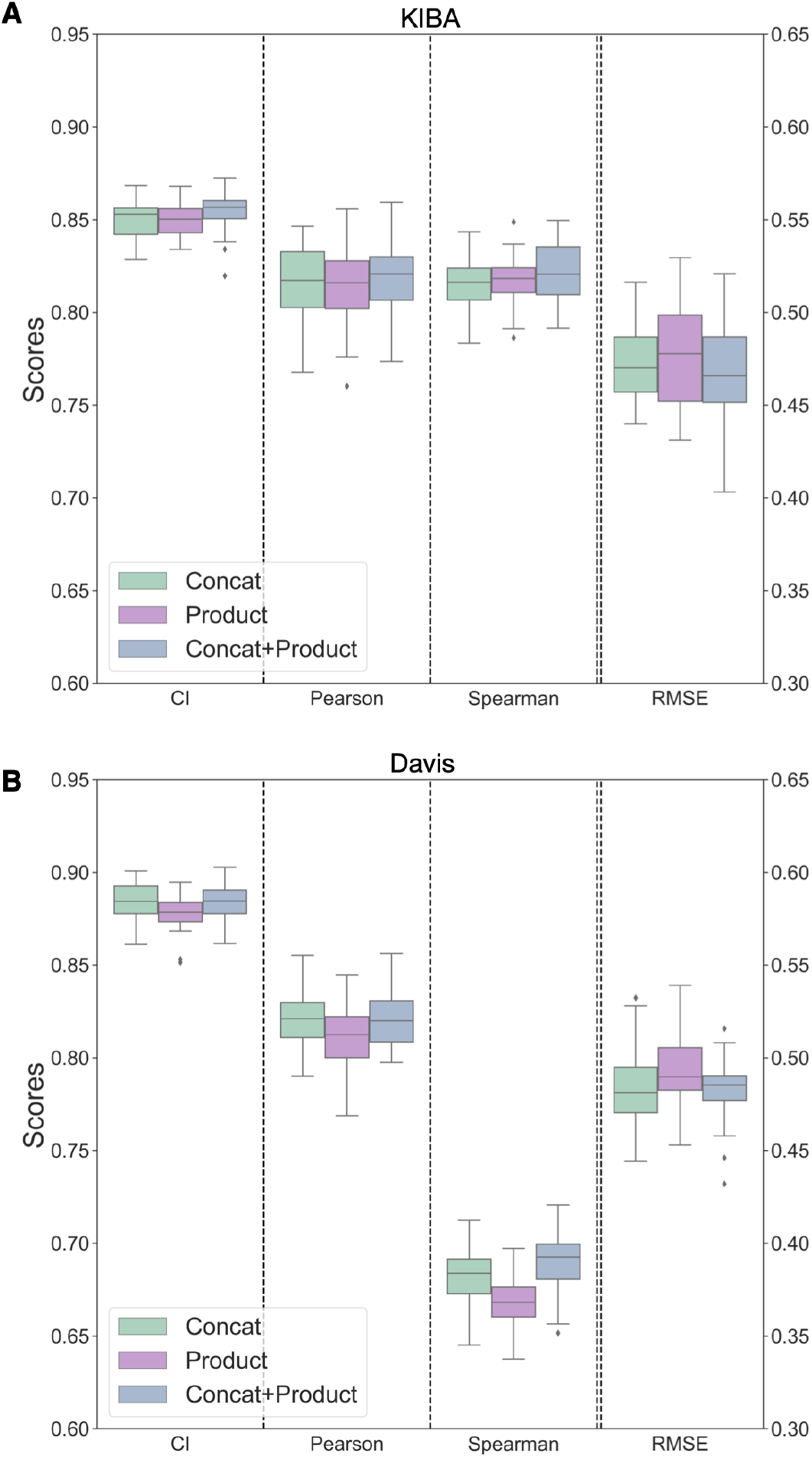
Performance of encoding combining techniques on both KIBA (A) and Davis (B) data sets show minimal change. Binding affinity prediction is minimally affected by the element-wise product and the concatenated vector obtained by combining both element-wise product and concatenation operations.

## Discussions and Conclusions

It is often most enticing for a new study to look at binding affinity predictions to introduce a new algorithm or machine learning model, which is then often superficially compared in performance (accuracy in terms of RMSE or correlation) to previous approaches. What is often neglected is looking at good comparison tools for assessing if a new model is statistically actually better than a previous model. What is often forgotten is that the training process is not deterministic, meaning we just pick the best-performing model after training but do not assess its variability. Here, we introduced robust significance testing and error analysis using Wilcoxon’s signed rank test to make sure we can make statistically significant statements when comparing our differently trained models on the same deep learning framework. We also included a robust bootstrapping error analysis often neglected when new models are introduced. All of this allowed us to carry out a detailed investigation in terms of how different parts of a deep learning model actually contribute to the overall performance of the final downstream tasks, i.e. the prediction of a binding affinity. In this paper, we systematically investigated the contributions of ligand and protein encodings to the downstream tasks of binding affinity predictions using 1D type of data and 2D type of data. Our 1D data encodings come from protein language models or hand-curated KLIFS data for proteins using convolutional neural network-based DL architectures. For the 2D data, we use graph-based approaches for both ligands, where SMILES strings get converted to ligand graphs and protein sequences get converted to a protein contact map either from a protein structure or through a protein language model. The deep learning CNN and GCN architectures used were not novel, however, we gained new insights into how protein and ligand embeddings contribute systematically to the learning of binding affinities for commonly used data sets used in the literature (KIBA and Davis). We successfully show that protein encodings, as often used in the literature [32], have little to no contribution to the downstream learning tasks, and all the correlation learned between structure and binding affinity is through ligand encodings. Furthermore, augmenting datasets from a 1D language model to a 2D graph model does not make the learning process significantly better. While we highlight the importance of understanding the role of encodings in DL models and provide insights into what the current DL models are learning in predicting protein-ligand binding affinity, we recognize other limitations associated with these DL methods. Most of the current DL methods are trained and tested on small-size kinase data sets with skewed binding interactions data (Figure 2). Testing DL frameworks only on kinase data sets is an obvious choice because of the amount of available data. However, care should be taken with the KIBA data set. It is tempting to augment a data set to include experimental data from multiple experimental assays to include *IC*_50_, *K_i_*, and *K_d_*. To broaden the dataset, the KIBA score was designed to account for multiple sources of experimental measurements [10]. Unfortunately, by combining information from different assays and measurement types the resulting uncertainty introduced is not accounted for in the KIBA score. This means evaluating the accuracy on the downstream task becomes inaccurate, as there is no notion of reliability of a single KIBA score incorporated into the model. As Killiokoski et al. [49] pointed out IC50s can be augmented with *K_i_*, but care should be taken when looking at SAR/QSAR models in terms of the maximally achievable performance due to the introduced noise. Here we chose to include KIBA results to compare to other existing methods but would recommend not using the data set as is in the future. More generally, the kinase data sets are a good starting point for model development but are not the most effective way to build a useful binding affinity prediction algorithm. Additionally, data sets with a more diverse set of proteins, such as BindingDB [50], will increase the generalizability to unseen proteins and ligands and test out new and better encodings.

We have seen that the DL models do not learn information that captures the protein and ligand interaction features but are biased toward learning from the ligand features. The current approaches to encode proteins from sequences and contact maps in the form of 1D and 2D encodings with CNN or GNN architectures are not sufficient to capture protein features to build a robust binding affinity prediction tool, and future work in this direction is required. There are exciting efforts making use of 3D protein-ligand interaction features from 3D complex structures [51, 52]. There is work to be done both in the area of 3D and 1-and 2D encodings to optimally learn protein-ligand interactions. One avenue to explore in the future is to jointly learnt features for making predictions; in this way, the DL model uses the joint features corresponding to protein-ligand interaction properties. This could be done, e.g., by including physics-based 3D snapshots in training the DL models to predict the binding affinity, similar to what has been explored to active learning approaches incorporating molecular dynamics-based binding affinity predictions [53].

## Acknowledgement

This work was supported by the United Kingdom Research and Innovation (grant EP/S02431X/1), UKRI Centre for Doctoral Training in Biomedical AI at the University of Edinburgh, School of Informatics and Exscientia Plc, Oxford. The authors thank John Chodera for his discussions and insights during the project.

