## Supplementary material for "From Proteins to Ligands: Decoding Deep Learning Methods for Binding Affinity Prediction": SI

Rohan Gorantla,[*,†,‡] Alžbeta Kubincová,[¶] Andrea Y. Weiße,[§,†] and Antonia S. J. S. Mey[*,‡]

†*School of Informatics, University of Edinburgh, EH8 9AB*

‡*EaStCHEM School of Chemistry, University of Edinburgh, EH9 3FJ*

¶*Exscientia, Schrödinger Building, Oxford, OX4 4GE*

§*School of Biological Sciences, University of Edinburgh, EH9 3FF*

### Supporting Information Available

#### Methods

##### Deep Learning model architectures and implementation details

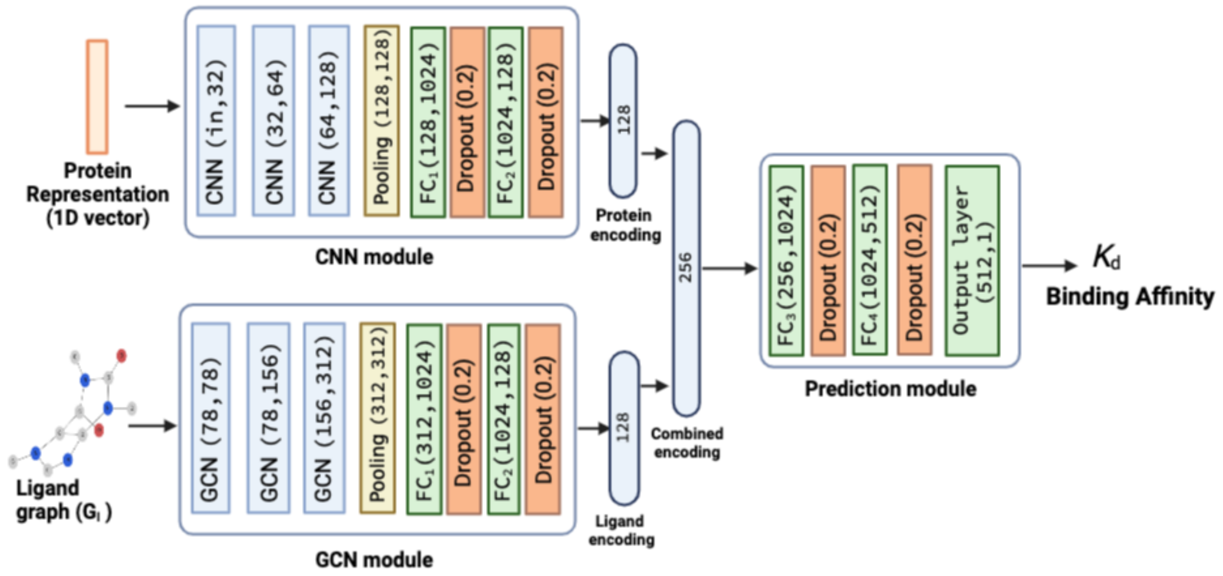


Figure S1: Convolutional neural network (CNN), GCN, and binding affinity prediction modules for studying the impact of 1D protein encodings and ligand graphs. The neural network layers in both the modules are shown along with their input and output channel sizes, i.e., *(input channel size, output channel size)*. The GCN module and prediction module architecture is the same as Figure S2. 1D protein representations (`in` is 1785 for KLIFS and 1280 for ESM) and ligand graphs are passed through the CNN module and GCN module respectively. For the proteins, features are first extracted from three 1D CNN layers with stride as 1, and increasing kernel sizes $[4, 8, 12]$. The output from the 1D CNN layers is then passed through the pooling layer and the pooling output is passed through two fully connected (FC) layers. The flattened feature vector of 128 dimensions is given as input to the first $FC_1$ layer in CNN module with 1024 neurons and outputs a feature vector of $1 \times 1024$ dimension. A dropout layer is added after the FC layer with a dropout rate of 0.2. The output from $FC_1$ layer is passed to $FC_2$ layer with 128 neurons. We obtain 128-dimensional protein and ligand encodings which are then combined to form a 256-dimensional embedding. We implement the 1D-CNN layer with `torch.nn.Conv1d()`, pooling layer with `torch.nn.AdaptiveMaxPool1d()`, FC and output layers with `torch.nn.Linear()` and the dropout layer with `torch.nn.Dropout()` class.

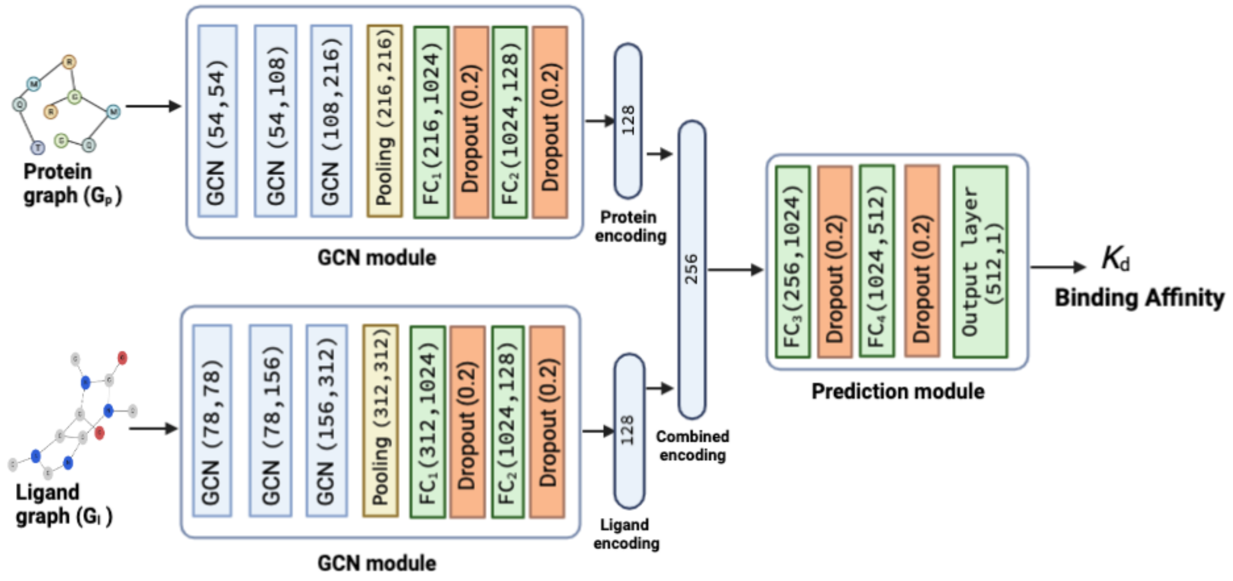


Figure S2: Graph convolutional network (GCN) and binding affinity prediction modules for studying the impact of protein and ligand graphs. The neural network layers in both the modules are shown along with their input and output channel sizes, i.e., *(input channel size, output channel size)*. Protein and ligand graphs are passed through the GCN module, here features are first extracted from 3 GCN layers and then passed through the pooling layer to get a single column vector of dimension **g** (**g** is 216 and 312 for protein and ligand graphs, respectively). These vectors from the pooling layer are then passed through two fully connected (FC) layers. The flattened feature vector of dimension **g** is given as input to the first $FC_1$ layer with 1024 neurons and outputs a feature vector of $1 \times 1024$ dimension. A dropout layer is added after the FC layer with a dropout rate of 0.2 to avoid overfitting and co-adaptation issues. The output from $FC_1$ layer is passed to $FC_2$ layer with 128 neurons. We obtain 128-dimensional protein and ligand encodings which are then combined to form a 256-dimensional embedding. We pass the 256-dimensional embedding into the prediction module with two FC layers and one output layer. The $FC_3$ and $FC_4$ layers have 1024 and 512 neurons, respectively, and a dropout layer is added after them. The output layer is similar to an FC layer with one neuron, and it takes 512-dimensional embedding to predict the binding affinity score. We implemented GCN layer with `torch_geometric.nn.GCNConv()`, pooling layer with `torch_geometric.nn.global_mean_pool()`, FC and output layers with `torch.nn.Linear()` and the dropout layer with `torch.nn.Dropout()` class. For studying element-wise product, we take an element-wise product of the 128-dimensional protein and ligand encoding to obtain the a 128-dimensional vector which is fed to the $FC_3$ layer instead of 256-dimensional vector in the case of concatenation. Similarly, for combined concatenation and element-wise product vector, the $FC_3$ layer will get an input vector of 384 dimensions.

#### Node features in a protein graph

Position-Specific Scoring Matrix (PSSM) provides the per-residue evolution patterns in the sequence profile [1]. By first counting the instances of each residue at each position, a position frequency matrix $\mathbf{M}^{\mathrm{PFM}}$ is generated using the equation below [2]

$$M_{\mathrm{k,j}}^{\mathrm{PFM}} = \sum_{i=1}^{Z} I\left(S_{i,j} = k\right), \tag{1}$$

where $S$ is a set of $Z$ aligned sequences for a protein sequence with length of $L_{\mathrm{p}}$, $k$ belongs to residue symbols set, $i = (1, 2, \ldots, Z)$, $j = (1, \ldots, L_{\mathrm{p}})$ and $I(x)$ is an indicator function when the condition $x$ is satisfied and 0 otherwise. A position probability matrix $\mathbf{M}^{\mathrm{PPM}}$ is then computed from the $\mathbf{M}^{\mathrm{PFM}}$ matrix using the following equation

$$M_{\mathrm{k,j}}^{\mathrm{PPM}} = \frac{M_{\mathrm{k,j}}^{\mathrm{PFM}} + \frac{c}{4}}{Z + c}, \tag{2}$$

where $c$ is the added pseudo count that is empirically set to 0.8 similar to [2] to avoid matrix entries with a value of 0 [3]. The $\mathbf{M}^{\mathrm{PPM}}$ matrix is then utilized to compute 21 PSSM features. For computing PSSM features, we need an aligned protein sequence in PSICOV [4] format. We used the aligned protein sequences in PSICOV format provided by Jiang et al. [2]. Table S1 below contains information about the rest of the node features.

##### KIBA metric

Kinase inhibitor bioactivity (KIBA) score is a continuous value of binding affinity developed by Jing et al. [5] integrating the information from biological activity of kinase inhibitors from $K_{\mathrm{i}}$, $\mathrm{IC}_{50}$, and $K_{\mathrm{d}}$ into a single bioactivity score [5]. Lower KIBA score denotes higher binding affinity. The KIBA score can be defined based on $K_{\mathrm{d}}$ or $K_{\mathrm{i}}$, or the average of them, depending on the availability of the bioactivity types [5] using the equation below

$$\begin{aligned} \mathrm{KIBA} &= \frac{\mathrm{IC}_{50}}{1 + H_{\mathrm{i}}\left(\mathrm{IC}_{50}/K_{\mathrm{i}}\right)} && \text{if } K_{\mathrm{i}} \text{ and } IC_{50} \text{ are present} \\ &= \frac{\mathrm{IC}_{50}}{1 + H_{\mathrm{d}}\left(\mathrm{IC}_{50}/K_{\mathrm{d}}\right)} && \text{if } K_{\mathrm{d}} \text{ and } IC_{50} \text{ are present} \\ &= \left(\frac{\mathrm{IC}_{50}}{1 + H_{\mathrm{i}}\left(\mathrm{IC}_{50}/K_{\mathrm{i}}\right)} + \frac{\mathrm{IC}_{50}}{1 + H_{\mathrm{d}}\left(\mathrm{IC}_{50}/K_{\mathrm{d}}\right)}\right)/2 && \text{if } K_{\mathrm{i}},\ K_{\mathrm{d}} \text{ and } IC_{50} \text{ are present,} \end{aligned} \tag{3}$$

Table S1: Residue node features in a protein graph

| **Node Features** | **Dimensions** |
|---|---|
| One-hot encoding of the residue symbol | 21 |
| Position-specific scoring matrix | 21 |
| Whether the residue is aromatic | 1 |
| Whether the residue is aliphatic | 1 |
| Whether the residue is polar neutral | 1 |
| Whether the residue is acidic charged | 1 |
| Whether the residue is basically charged | 1 |
| Residue weight | 1 |
| Negative of the logarithm of the dissociation constant for the −COOH group | 1 |
| Negative of the logarithm of the dissociation constant for the −NH group | 1 |
| Negative of the logarithm of the dissociation constant for any other group in the molecule | 1 |
| pH at the isoelectric point | 1 |
| Hydrophobicity of residue (pH = 2) | 1 |
| Hydrophobicity of residue (pH = 7) | 1 |
| *Total* | 54 |

where $H_i$ and $H_d$ are the parameters that determine the weights of $IC_{50}$ in the model-based adjustments for $K_i$ and $K_d$.

#### Results

##### Contact map evaluation

The Matthews Correlation Coefficient (MCC) [6] is a widely used metric to assess the quality of binary classifiers, including those used in protein contact prediction. In this context, MCC measures the agreement between predicted and true contact maps, which are binary matrices indicating the presence or absence of contact between pairs of residues in a protein. MCC can be calculated from a confusion matrix that summarizes the number of true positives (TP), true negatives (TN), false positives (FP), and false negatives (FN) obtained by comparing the predicted contact map to the true contact map. The MCC is given by

$$\mathrm{MCC} = \frac{\mathrm{TP} \times \mathrm{TN} - \mathrm{FP} \times \mathrm{FN}}{\sqrt{(\mathrm{TP}+\mathrm{FP}) \times (\mathrm{TP}+\mathrm{FN}) \times (\mathrm{TN}+\mathrm{FP}) \times (\mathrm{TN}+\mathrm{FN})}} \tag{4}$$

**Algorithm 1:** Point Randomization of Ligands

**Input** : `SMILES` string, $S_i$

**Initialization:** Get the count of Cl, F, Br, C and (=O) atoms from the input $S_i$ and set to #Cl, #F, #Br, #C and #(=O) respectively.

```
N ← #Cl + #F + #Br + #(=O);
if N > 0 AND N ≤ 4 then
    Randomly select n changes to be made from the range of 1 to N possible
      changes;
    while n ≠ 0 do
        Randomly select an atom from Cl, F, Br, and (=O) which are present in S_i;
        if Halogen (i.e., Cl) is selected then
            Replace selected halogen (Cl) from S_i with one of other halogens (F or
              Br) randomly to obtain an updated SMILES string S_i ;
            #Cl ← #Cl − 1 ;                    /* Update the Halogen count */
            n ← n − 1;
            continue
        end
        if (=O) is selected then
            Remove one (=O) from S_i to obtain an updated SMILES string S_i ;
            #(=O) ← #(=O) − 1;
            n ← n − 1;
            continue
        end
    end
else
    S_i ← Cl+ S_i ;
end
```

**Output:** Updated `SMILES` string, $S_o$

MCC values range from -1 to 1 , with 1 indicating perfect agreement, 0 indicating random prediction, and -1 indicating complete disagreement between predicted and true contact maps. One advantage of using MCC in cases with imbalanced datasets is that it takes into account both true positives and true negatives, as well as false positives and false negatives, to provide a balanced assessment of the performance of the classifier. This is particularly important when the positive and negative classes are not balanced in the dataset, as metrics such as accuracy can be misleading. In the context of contact map prediction, the true contacts (positive class) are typically much rarer than the non-contacts (negative class), resulting in an imbalanced dataset.

Table S2: Overview of various experiments performed in our study

| Protein encoding | Ligand encoding | Combining method |
|---|---|---|
| *To study the impact of 1D and 2D protein encodings* | | |
| Pconcs4 | Original ligand graph | Concatenation |
| Alphafold2 | Original ligand graph | Concatentation |
| ESM | Original ligand graph | Concatentation |
| Random | Original ligand graph | Concatentation |
| KLIFS (1D) | Original ligand graph | Concatentation |
| ESM (1D) | Original ligand graph | Concatentation |
| *To study the impact of ligand encodings* | | |
| Pconcs4 | Point randomisation | Concatentation |
| Pconcs4 | Random node features | Concatentation |
| Pconcs4 | Random sampling | Concatentation |
| *Assessing the impact of combining methods* | | |
| Pconcs4 | Original ligand graph | Element-wise product |
| Pconcs4 | Original ligand graph | Concatenation + Element-wise product |

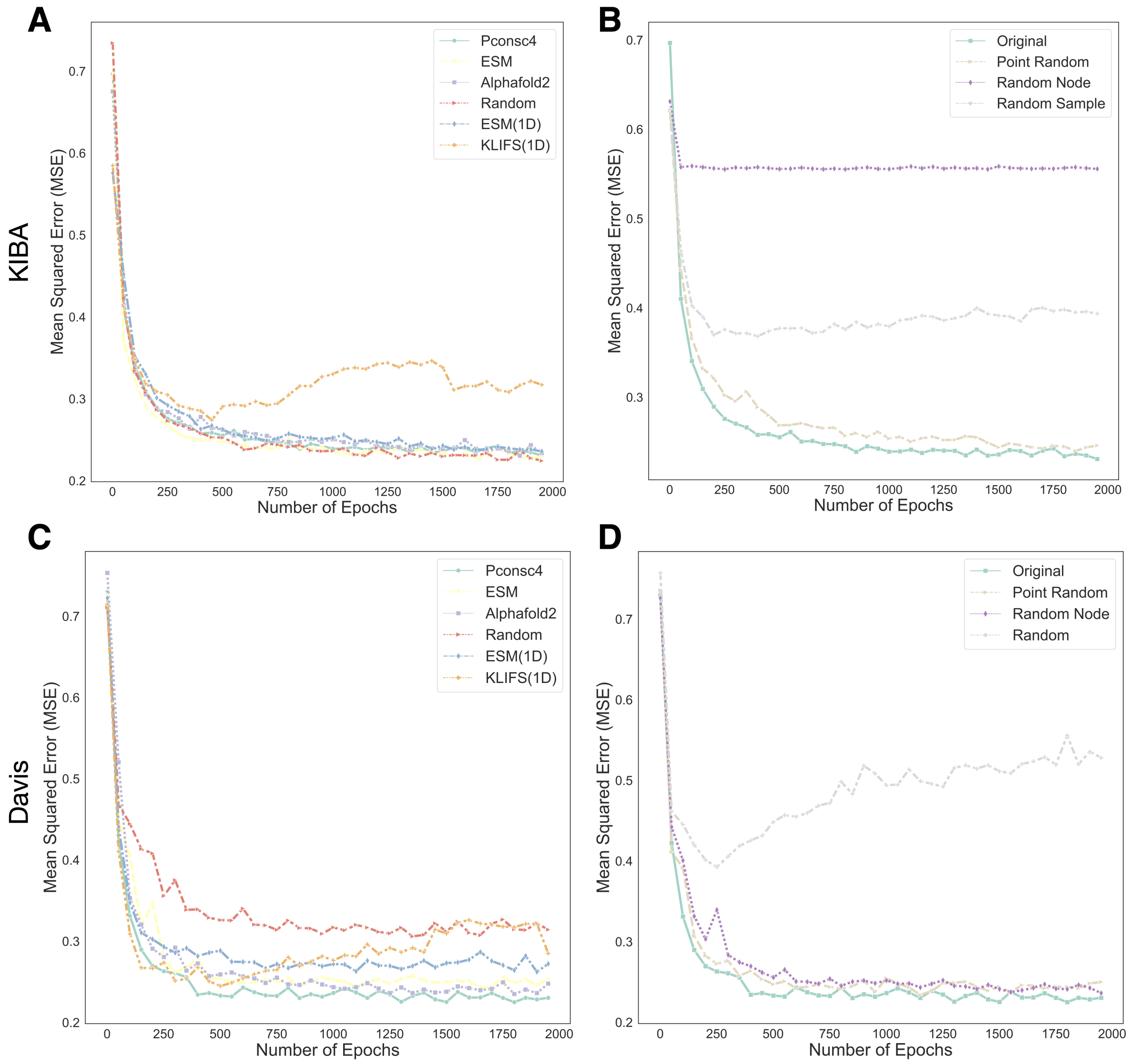


Figure S3: Mean squared error (MSE) curves on validation data during the training of DL models using various protein and ligand encodings. **A, C** MSE curves for protein encodings on the KIBA and Davis dataset. **B, D** MSE curves for ligand encodings on the KIBA and Davis dataset.

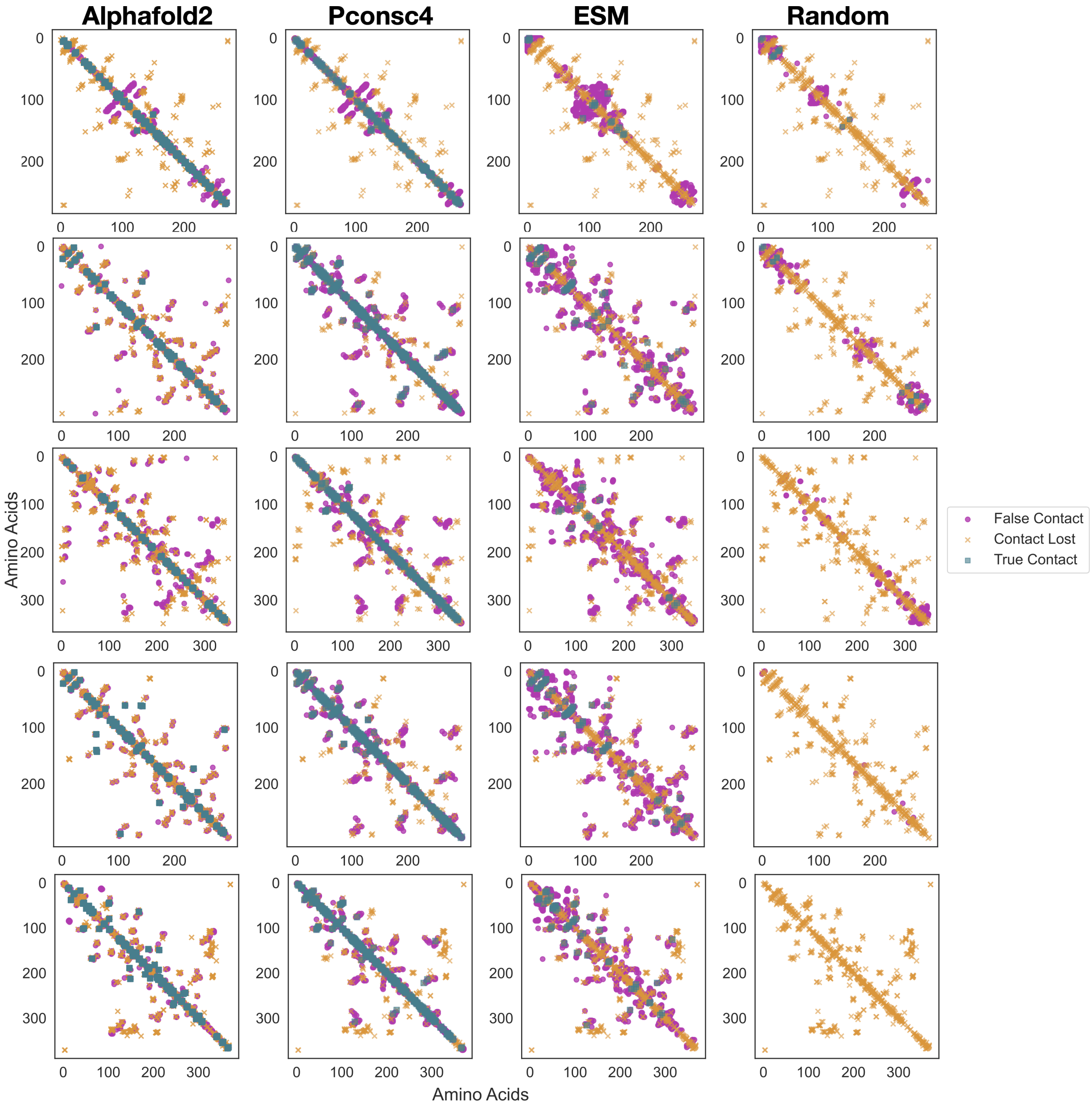


Figure S4: Visual illustration of contact maps obtained from various kinases present in KIBA and Davis datasets as compared to the contact maps obtained from PDB structures of kinases. The true contacts, false contact and contacts lost are highlighted.

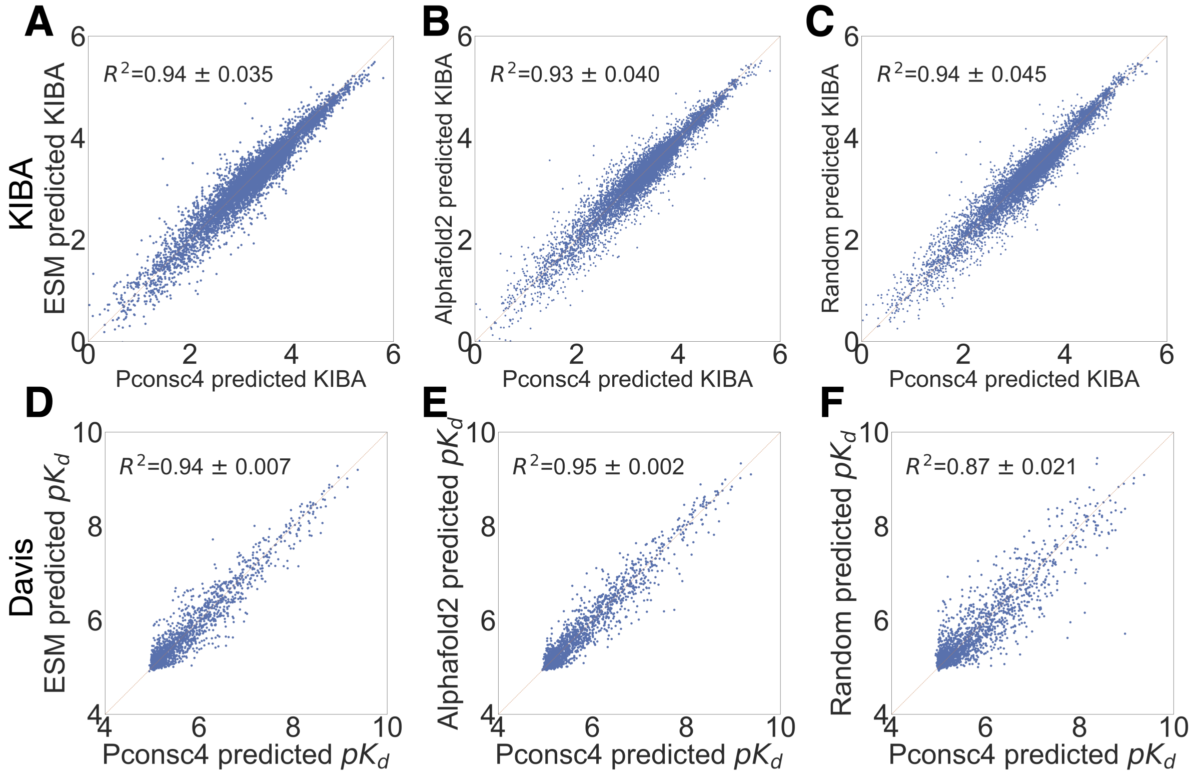


Figure S5: Correlation between binding affinity predictions given by the graph-DL model with different protein encoding methods- **A, B, and C:** KIBA and **D, E, and F:** Davis. The protein encoding based on random contact map is also strongly correlated with Pconsc4 methods. All these scatter plots show that the BA predictions are not impacted by the protein encodings obtained from various contact map methods and function in the same way on both the datasets.

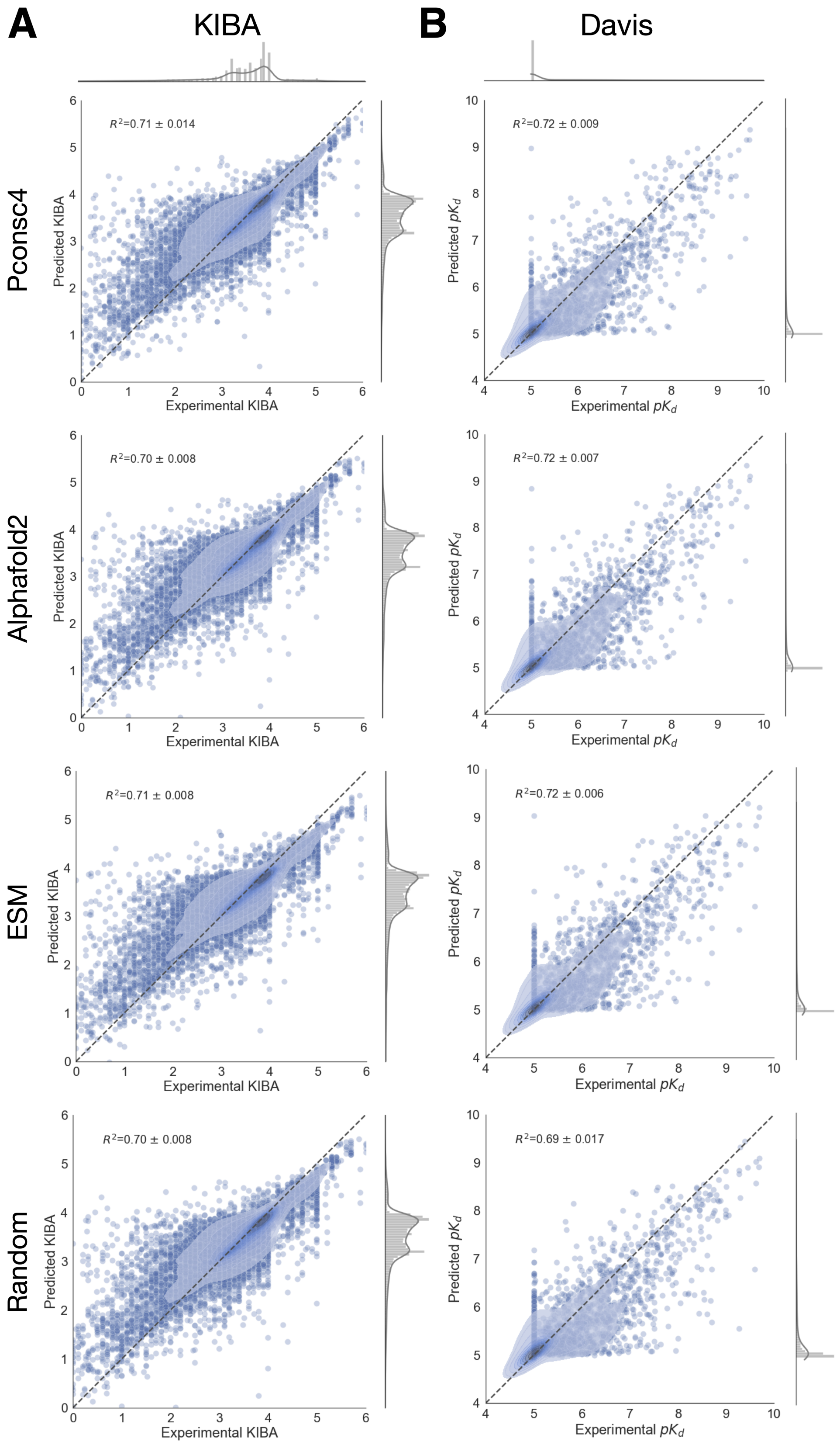


Figure S6: Experimental vs. predicted binding affinities of DL model trained using four different 2D encodings generated from contact maps on both KIBA and Davis datasets.

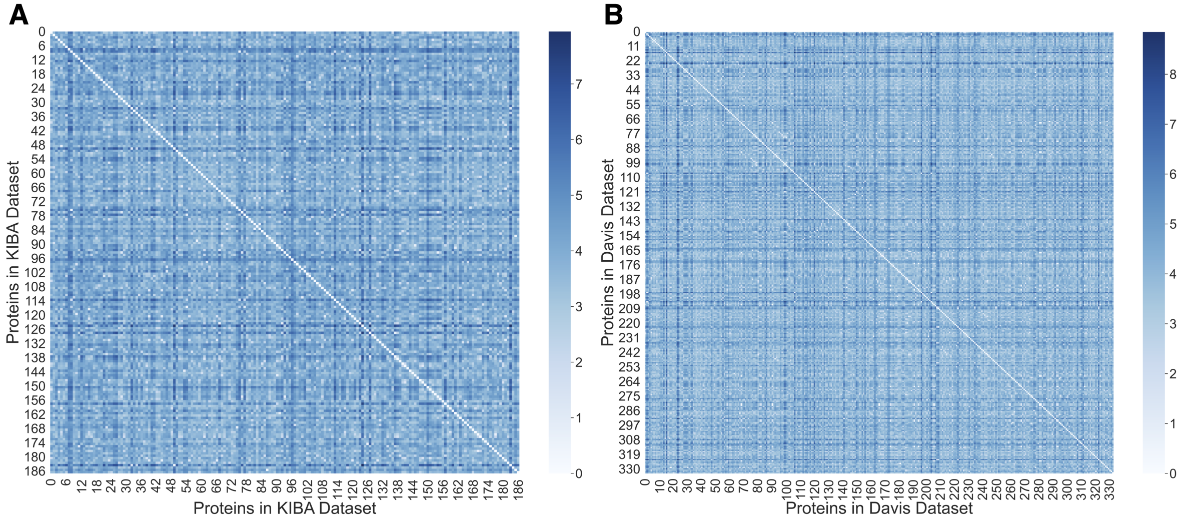


Figure S7: Euclidean distance between the ESM embeddings of proteins in the Davis and KIBA datasets. These embeddings capture 95% variance among the kinases in the Davis and KIBA datasets, and from the heatmap we can see that the Euclidean distance between the ESM embeddings of kinases is considerable, indicating a substantial distance between them in the Euclidean space. **A:** ESM embeddings from 188 KIBA proteins. **B:** ESM embeddings from 334 Davis proteins.

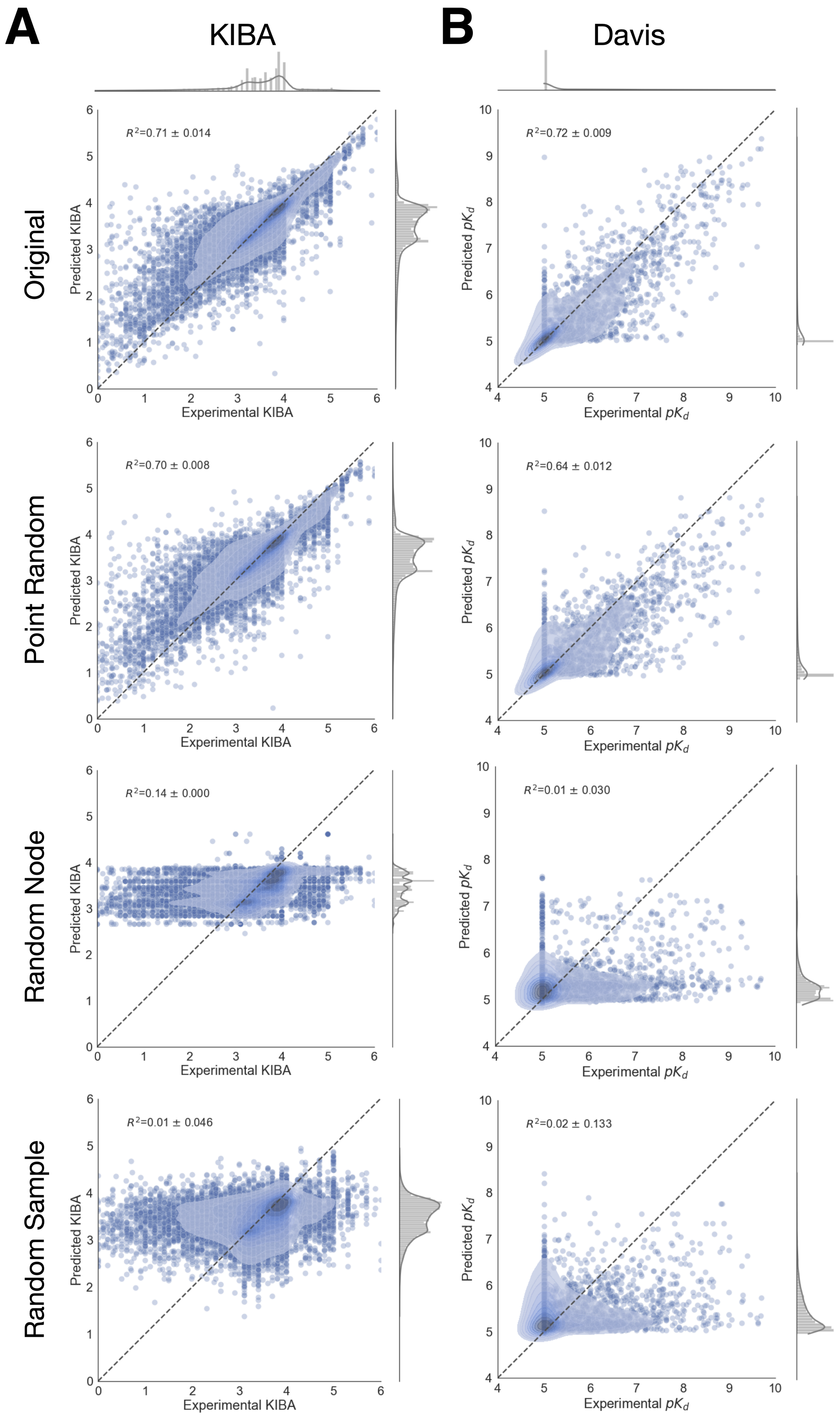


Figure S8: Experimental vs. predicted binding affinities of DL model trained using various perturbations of ligand graph encodings on both KIBA and Davis datasets.
